# mRNA nuclear clustering leads to a difference in mutant huntingtin mRNA and protein silencing by siRNAs *in vivo*

**DOI:** 10.1101/2024.04.24.590997

**Authors:** Sarah Allen, Daniel O’Reilly, Rachael Miller, Ellen Sapp, Ashley Summers, Joseph Paquette, Dimas Echeverria Moreno, Brianna Bramato, Nicholas McHugh, Ken Yamada, Neil Aronin, Marian DiFiglia, Anastasia Khvorova

## Abstract

Huntington’s disease (HD) is an autosomal dominant neurodegenerative disease caused by CAG repeat expansion in the first exon of the huntingtin gene (*HTT*). Oligonucleotide therapeutics, such as short interfering RNA (siRNA), reduce levels of huntingtin mRNA and protein *in vivo* and are considered a viable therapeutic strategy. However, the extent to which they silence HTT mRNA in the nucleus is not established. We synthesized siRNA cross-reactive to mouse (wild-type) *Htt* and human (mutant) *HTT* in a di-valent scaffold and delivered to two mouse models of HD. In both models, di-valent siRNA sustained lowering of wild-type *Htt*, but not mutant *HTT* mRNA expression in striatum and cortex. Near-complete silencing of both mutant HTT protein and wild-type Htt protein was observed in both models. Subsequent fluorescent in situ hybridization (FISH) analysis shows that di-valent siRNA acts predominantly on cytoplasmic mutant *HTT* transcripts, leaving clustered mutant *HTT* transcripts in the nucleus largely intact in treated HD mouse brains. The observed differences between mRNA and protein levels, exaggerated in the case of extended repeats, might apply to other repeat-associated neurological disorders.

## Introduction

Huntington’s Disease (HD) is a rare autosomal dominant neurodegenerative disease that causes progressive motor and cognitive impairments, eventually leading to death (1). No disease-modifying treatments are available.

HD is caused by a CAG repeat expansion in exon 1 of the *huntingtin* gene (*HTT*), resulting in transcription of mutant *HTT* mRNA with an expanded CAG tract and translation of mutant HTT protein with an expanded polyglutamine (polyQ) tract (2). The role of mutant HTT protein has been the primary focus of investigations into HD pathogenesis, with multiple phenotypes being reported (3). Recent studies indicate that uninterrupted CAG repeat length, not polyQ length (encoded by CAA-CAG), affects the rate of pathogenesis in HD patients, suggesting the potential importance of mutant *HTT* DNA*/*mRNA in HD (4). Indeed, in mouse models of HD, introducing a copy of *huntingtin* containing the mutant expanded CAG tract leads to HD-like phenotypes *in vivo* (5–7). The ability of different compounds to modulate not only the protein but also mutant *HTT* mRNA needs to be considered.

The presence of the expanded CAG repeat tract has an impact on the intracellular localization of *HTT* mRNA. Whereas wild-type *Htt* is predominantly cytoplasmic, mutant *HTT* mRNA is retained in the nucleus and forms insoluble clusters (7, 8). AGO2, the protein mediating small interfering RNAs (siRNA) activity, has been historically considered a cytoplasmic protein though it can be localized to the nucleus (9). The impact of localization on siRNA efficacy has not yet been examined in *HTT* mRNA.

Oligonucleotide therapeutics, including siRNAs and antisense oligonucleotides (ASOs), enable sequence-specific lowering of gene expression and are being explored as potential therapeutics for HD (10, 11). siRNAs harness endogenous RNA interference (RNAi) to enable targeted gene silencing at the mRNA level, thereby preventing translation of the corresponding protein in the cytoplasm (12). We recently developed an siRNA scaffold for delivery to the central nervous system (CNS), termed di-valent siRNA, which supports broad distribution and modulation of target gene expression in mouse and non-human primate CNS for up to six months following intracerebroventricular (ICV) injection (13). When programmed with an siRNA sequence complementary to both wild-type *Htt* and mutant *HTT*, di-valent siRNA silenced wild-type Htt protein and mutant HTT protein with similar efficacy in multiple HD mouse modes (14, 15).

Here, during evaluation of the ability of di-valent siRNA to silence wild-type *Htt* mRNA and mutant *HTT* mRNA in two transgenic mouse models of HD we discovered a profound difference in observed remaining levels of mRNA and protein. YAC128 and BAC-CAG express wild-type mouse *Htt* (7 CAG repeats) and an artificial copy of the human mutant *HTT* gene (128 mixed CAA/CAG repeats for YAC128, 81 uninterrupted CAG repeats; 120 CAG repeats for BAC-CAG) (5, 6). Di-valent siRNA induced potent silencing of wild-type *Htt*, but not mutant *HTT*, mRNA. Nevertheless, near-complete silencing of both wild-type Htt and mutant HTT protein was observed. Fluorescent in situ hybridization (FISH) analysis using RNAscope showed that di-valent siRNA silenced cytoplasmic wild-type Htt mRNA and mutant HTT mRNA – explaining the near-complete silencing of both mutant and wild-type HTT protein – but had minimal effect on nuclear mutant *HTT* mRNA clusters. Our findings reveal that discrepancies in siRNA mediated silencing of mRNA and protein may be explained by the localization and structure of target mRNA transcripts.

## Materials and Methods

### Oligonucleotides

Oligonucleotides were synthesized on a MerMade12 (Biosearch Technologies) using standard solid-phase phosphoramidites. Phosphoramidites used include: 2’-Fluoro RNA or 2’-*O*-Methyl RNA modified phosphoramidites (ChemGenes and Hongene Biotech), 5’-(*E*)-Vinyl tetra phosphonate (pivaloyloxymethyl) (Hongene), and 2’-O-methyl-uridine 3’-CE phosphoramidite (Hongene). Phosphoramidites were prepared at 0.1 M in anhydrous acetonitrile (ACN), except for 2’-O-methyl-uridine phosphoramidite which was dissolved in anhydrous ACN containing 15% dimethylformamide. 5-(Benzylthio)-1H-tetrazole (BTT) activator was used at 0.25 M. Coupling time and excess phosphoramidite were 4 mins and 10 eq, respectively. All synthesis reagents were purchased from Chemgenes, unless otherwise specified. Unconjugated oligonucleotides were synthesized on 500 Å long-chain alkyl amine (LCAA) controlled pore glass (CPG) functionalized with Unylinker terminus. Divalent oligonucleotides (DIO) were synthesized on a custom solid support prepared in-house (13).

Vinyl-phosphonate-containing oligonucleotides were deprotected with 3% diethylamine in ammonium hydroxide, for 20 hours at 35°C with agitation. Di-valent oligonucleotides were deprotected with 1:1 ammonium hydroxide and aqueous methylamine for 2 h at 25°C with agitation. The products were dried overnight before being filtered and rinsed with 15 mL of 5% ACN in water. Purifications were performed on an Agilent 1290 Infinity II HPLC system using Source 15Q anion exchange resin (Cytiva, Marlborough, MA). The loading solution was 20 mM sodium acetate in 10% ACN in water, and elution solution was the loading solution with 1 M sodium bromide, using a linear gradient from 30% to 70% in 40 mins at 50°C. Pure fractions were combined and desalted by size-exclusion chromatography with Sephadex G-25 (Cytiva). Oligonucleotide purities and molecular weights were confirmed by IP-RP LC-MS on an Agilent 6530 Accurate-mass Q-TOF LC-MS.

Single strands were annealed by adding a 1:1 ratio of antisense strand to sense strand in water at the desired concentration. The mixture was heated at 95°C for 10-15 mins, then removed and allowed to cool at room temperature. A 20% TBE gel (Invitrogen, EC63152BOX) was used to verify successful annealing. siRNAs were dried down using a speed vacuum concentrator, then resuspended in calcium and magnesium-enriched artificial CSF (137 mM NaCl, 5 mM KCl, 14 mM CaCl_2_, 2 mM MgCl_2_, 20 mM glucose, 8 mM HEPES, the pH was adjusted to 7.4 with NaOH). Di-siRNA were dosed to 440 μg (20 nmol) per mouse.

### Animal studies

All mouse studies were approved by the UMass Chan IACUC (Protocol # 202000010) and completed in accordance with the National Institutes of Health Guideline for Laboratory Animals. BAC-CAG and YAC128 heterogenous FVB mice were obtained in equal ratios of males to females from the Jackson Laboratory. No sex differences were observed. Animals were housed at pathogen-free animal facilities at UMass Chan Medical School on a 12-hour light/12-hour dark cycle at a controlled temperature (23 ± 1 °C) with standard humidity (50% ± 20%). Animals were maintained in cages holding a maximum of five mice of either sex and were given access to food and water *ad libitum*. YAC128 male mice were bred with wild-type FVB female mice, resulting in mixed wild-type and heterogenous litters. Litters were genotyped by performing PCR on DNA extracted from ear punches taken during weaning.

### In vivo stereotactic injection of oligonucleotides

Adult YAC128 and BAC-CAG mice were 8 and 12 weeks old, respectively, at the time of injection. Prior to injection, mice were anesthetized with (2.5%) vaporized isoflurane. 10 nmol of di-siRNA was administered into each ventricle at a rate of 1 μL per minute. All injections were performed using sterile surgical techniques and used a standard rodent stereotaxic instrument and an automated microinjection syringe pump (Digital Mouse Stereotaxic Frame; World Precision Instrument #504926). A burr was used to drill a small hole in the following coordinates: −0.2 mm posterior, 1.0 mm mediolaterally. The needle was then placed at a depth of −2.5 mm ventrally. A volume of 5 μL was injected per ventricle at a rate of 750 nL/min. Following injection, mice were monitored until sternal. Mice were euthanized according to the UMass Chan IACUC protocol (#202000010). At the time of euthanasia (24 weeks post-injection for YAC128 mice, 4 weeks post-injection for BAC-CAG mice), mice were anesthetized with isoflurane and perfused intracardially with 20 ml 1X PBS buffer.

### Tissue processing

Brains were dissected out and bisected along the longitudinal fissure. The right side of the brain was sliced using a mouse brain-slicing matrix. 1.5 mm^3^ punches were taken from brain slices in the medial cortex and striatum. Punches for protein were flash-frozen on dry ice and stored at −80°C. Punches for mRNA were stored at 4°C in 100 μL of RNA later overnight or until use. mRNA brain tissue punches were homogenized in 96-well plate format on a QIAGEN TissueLyser II (Qiagen, Valencia, CA; #85300) in 600 μL of homogenizing buffer (ThermoFisher Scientific; #10642) and 10 μL of proteinase K (AM2546). For FISH analysis, the left sides of the brains were placed (anterior portion facing downwards) in disposable cryomold (Polysciences Inc. #18986– 1), and frozen in O.C.T. embedding medium (Tissue-Tek #4583) in a dry ice/ethanol bath. Brains were stored at −80°C until use and transferred overnight at −20°C prior to sectioning. Brains were sliced into 20-μm sections using a cryostat (temperatures: sample holder −13°C, blade −12°C) (ThermoFisher CryoStar™ NX70) and mounted on superfrost slides (Fisher #1255015). Sectioning occurred from the anterior end back, trimming 1500 μm then taking 20 μm sections. Six sections (two slides) were collected, then 200 μm was discarded, and another six sections were taken. This process was repeated for a total of 10 slides, which were stored at −80°C until use.

### Fluorescent in situ hybridization

Mouse brain sections were fixed in 10% formalin for one hour at 4°C and dehydrated in accordance with the manufacturer protocol for fresh frozen tissue (ACDBio #323100). FISH was performed using the RNAscope® Fluorescent V2 Multiplex kit (ACDBio #323100) following the manufacturer’s instruction (ACDBio #323100). Following sample preparation, samples were incubated with a combination of target probes against human huntingtin (ACDBio # 420231) and mouse huntingtin (ACDBio # 473001C2) in the HybEZ™ oven at 40°C for 2 hours, again in accordance with the RNAscope® Fluorescent V2 Multiplex kit manual (ACDBio #323100). Detection fluorophores used were Alexa fluor 488 (Thermofisher # B40953) for human HTT and Alexa fluor 647 (Thermofisher # B40958) for mouse Htt. Sample nuclei were stained with DAPI solution for 5 minutes, washed twice with PBS, mounted in ProLong™ Gold antifade medium (ThermoFisher #P36930), and dried at room temperature overnight before being sealed with clear nail polish.

### Microscopy and Image processing

Images were acquired using an Andor Dragonfly 505 on a Leica DMi8 using the confocal system with the Leica HC PL APO 63x/1.40 OIL CS2 oil objective and a Andor Zyla sCMOS 4.2 Plus (2x). Fusion (v2.4.0.14) software was used for image acquisition and z stacks were collected using the automatic step size function. Images were focused in the z plane using the ASI XY stage with a 500 um piezo. Image acquisition settings were kept consistent for all experiments and empirically determined using the manufacturer’s 3plex positive control probe set (ACDBio #320881) and 3plex negative control probe set (ACDBio #320871). Andor Integrated Laser Engine with 405/488/640 nm lasers were set to the following exposure times: 405 nm laser, 200 ms integration; 488 nm laser, 1000 ms; 640 nm laser, 1000 ms., and made use of the Triple emission filter 445-521-594.

Image processing of RNAscope images was performed with ImageJ using a macro written in-house (accessible at https://github.com/socheataly/imagej). The method of image processing and RNA foci quantification was done in accordance with the protocol described in Ly et al., 2022 (7). Parameters were kept consistent between all sample groups and were determined by using the manufacturer’s 3plex positive control and 3plex negative control probe sets such that no processed RNAscope signal was detectable in the 3plex negative control sample.

### mRNA and protein measurements

mRNA was quantified using the QuantiGene 2.0 Assay (ThermoFisher Scientific; #QS0016, QS0106). For tissue lysate, 40 μl of each lysate was added to the capture plate. Probe sets were diluted as specified in the Quantigene singleplex protocol and 20 μl human *HTT*, mouse *Htt*, or *Hprt* probe set (ThermoFisher Scientific; #SA-50339, #SA-10003) was added to appropriate wells for a final volume of 100 μl. Signal was amplified according to the Quantigene protocol. Luminescence was detected on either a Veritas Luminometer (Promega, #998–9100) or a Tecan M1000 (Tecan).

Protein quantification was performed by capillary immunoassay analysis using the automated simple western system, Wes (ProteinSimple). Frozen tissues from striatum or medial cortex were homogenized in 75 μL 10 mM HEPES (pH 7.2) with 250 mM sucrose, 1 mM EDTA + protease inhibitor tablet (Roche #04693116001; complete, EDTA-free) + 1 mM NaF +1 mM Na_3_VO_4_ and sonicated for 10 seconds at 4°C. Bradford assay was used to quantify total protein concentration. 0.6 μg of protein were analyzed with the Wes system following the manufacturer’s protocol with the following antibodies: huntingtin (1:50, Ab1, aa1-17, (19)), and Vinculin (1:5,000, Sigma, V9131). The peak area of each protein was automatically determined using Compass software (ProteinSimple) with the “dropped line” option on the electropherogram data and the level of HTT was normalized to the vinculin loading control. Images in Supplementary Figures 2-4 are shown as the “lane view” option of the electropherogram data.

### Quantification and statistical analysis

No statistical methods were performed to pre-determine sample sizes. *In vivo* studies used a starting n of six mice per group, while FISH analyses used an n of three mice. Data analyses were performed using GraphPad Prism 8.4.3 software (GraphPad Software, LLC) and an R script using the lme4 and emmeans packages (16, 17). For FISH measurements, in each sample group, at least 100 cells were analyzed. Statistical tests for FISH analyses were performed using a mixed effects model and analyzed using pairwise comparisons contrasting the impact of siRNA treatment. For mRNA and protein knockdown measurements, a one-way ANOVA with multiple comparisons was used to compare all groups to NTC. The statistical test used is reported in the figure legends with the level of statistical significance, which is denoted by asterisks (*, P < 0.05; **, P < 0.01; ***, P < 0.001; ****, P < 0.0001).

## Results

### Design of di-valent siRNA targeting wild-type *Htt* and mutant *HTT*

We synthesized a validated siRNA sequence cross-reactive to wild-type *Htt* and mutant *HTT* or a non-targeting control (NTC) sequence in the di-valent scaffold (13) with an optimized 2’-O-Methyl RNA/2’-Fluoro RNA pattern and phosphorothioated terminal backbones. These chemical modifications improve the potency and stability of di-valent siRNA *in vivo* (18, 19) (Supplementary Figure 1). A 5’ vinylphosphonate (5’VP) was used to improve phosphatase stability and improve silencing activity (20). Finally, we incorporated two extended nucleic acid (exNA) modifications at the 3’ end of the antisense strand to improve efficacy and nuclease resistance (21).

### Di-valent siRNA treatment reduces wild-type and mutant *HTT* mRNA with significantly different efficiencies in BAC-CAG mice

440 μg (20 nmol) of di-valent siRNA targeting *Htt*/*HTT* or the NTC control were delivered to 16-week-old BAC-CAG mice (n=6) via bilateral ICV injection (Figure 1A). Mice were sacrificed one-month post-injection. Wild-type and mutant *HTT* mRNA and protein levels were evaluated by QuantiGene and automated simple western, in the cortex and striatum (Figure 1B-C and Supplementary Figure 2). Di-valent siRNA achieved ∼65% silencing of wild-type *Htt* mRNA in both striatum (p=0.0026) and cortex (p=0.001) compared to NTC control. The reduction in mutant *HTT* mRNA compared to NTC control was not statistically significant in the striatum (p=0.1762) or cortex (p=0.0866). Near-complete silencing of HTT protein was observed in striatum and cortex for both mutant HTT (∼90% silencing, p=0.0144 in striatum; ∼95% silencing, p=0.0004 in cortex) and wild-type Htt (∼80% silencing, p=0.0313 in striatum; ∼85% silencing, p=0.0006 in cortex) protein. This discrepancy in degree of mRNA silencing versus protein silencing was most evident for mutant HTT (difference in residual mutant mRNA vs. protein is 58% in striatum and 55% in cortex).

**Figure 1:**
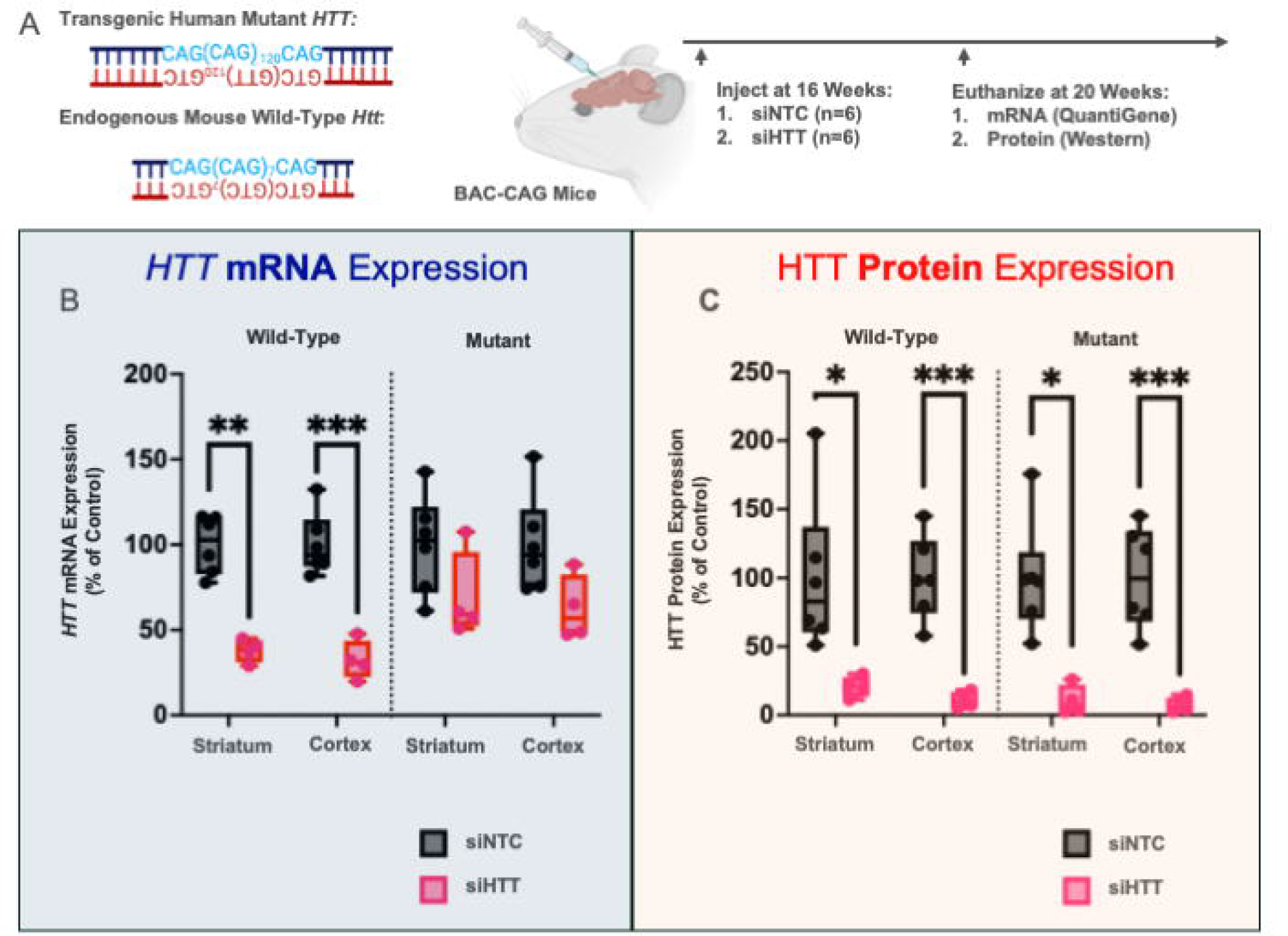
*HTT* mutant and wild-type mRNA but not protein are differentially affected by di-siRNA treatment in BAC-CAG mice after 1 month. (A) Three-month-old BAC-CAG mice were injected ICV with 10 μL of 20 nmol di-siRNA targeting either *HTT* or a NTC (n=6). BAC-CAG mice express both transgenic human mutant *HTT* with 120 CAG repeats and endogenous mouse wild-type *Htt* with 7 CAG repeats. Mice were sacrificed one month later at 20 weeks old. After sacrifice striatum and cortical tissue were tested for changes *HTT* mRNA (Quantigene) and protein (automated simple western) expression. (B) Human mutant *HTT* and wild-type mouse *Htt* mRNA levels were measured by Quantigene and normalized to *Hprt* as a housekeeping gene. mRNA levels are displayed as a percentage of their respective non-targeting control value. For comparisons between groups a one-way anova was used with multiple comparisons. For comparisons between siNTC and siHTT treated tissue p = 0.0026 and p= 0.001 for wild type *Htt* in the striatum and cortex respectively. For mutant *HTT*, p = 0.1762 and p = 0.0866 in the striatum and cortex respectively. (C) Human mutant HTT and wild-type mouse Htt protein levels were measured by automated simple western and normalized to vinculin as a loading control. Protein levels are displayed as a percentage of their respective non-targeting control value. For comparisons between groups a one-way anova was used with multiple comparisons. For comparisons between siNTC and siHTT treated tissue, p = 0.0313 and p= 0.0006 for wild type Htt in the striatum and cortex respectively. For mutant HTT p = 0.0144 and p = 0.0004 in the striatum and cortex respectively.

### Di-valent siRNA treatment reduces wild-type and mutant *HTT* mRNA with significantly different efficiencies in YAC128 mice

To evaluate if the observed difference in levels of *Htt* and *HTT* mRNA silencing is specific to the BAC-CAG model, we repeated the prior experiment in the YAC128 transgenic mouse model of HD (Figure 2A). 240 μg (10 nmol) of di-valent siRNA targeting *Htt/HTT* or the NTC control was delivered to 4-week-old YAC128 mice (n=6) via bilateral ICV injection. Mice were sacrificed six months post-injection, and wild-type Htt and mutant HTT mRNA and protein levels were measured in cortex and striatum as previously described (Figure 2B-C and Supplementary Figures 3 and 4). Di-valent siRNA achieved ∼50% silencing of wild-type *Htt* mRNA in both the striatum (p<0.0001) and cortex (p=0.0172) compared to NTC control. Similar to results in BAC-CAG mice, reduction in mutant *HTT* mRNA compared to NTC control did not reach statistical significance in cortex (∼30% reduction, p=0.3255). However, di-valent siRNA did achieve 25% silencing of mutant *HTT* mRNA in striatum (p= 0.0253) compared to NTC control. 85% silencing of mutant HTT (p<0.0001) and 90% silencing of wild-type Htt (p<0.0001) protein was observed. As in BAC-CAG mice, the difference in degree of mRNA versus protein silencing was more pronounced for mutant HTT (difference in residual mutant mRNA vs. protein is 63% in striatum and 58% in cortex) than for wild-type Htt.

**Figure 2:**
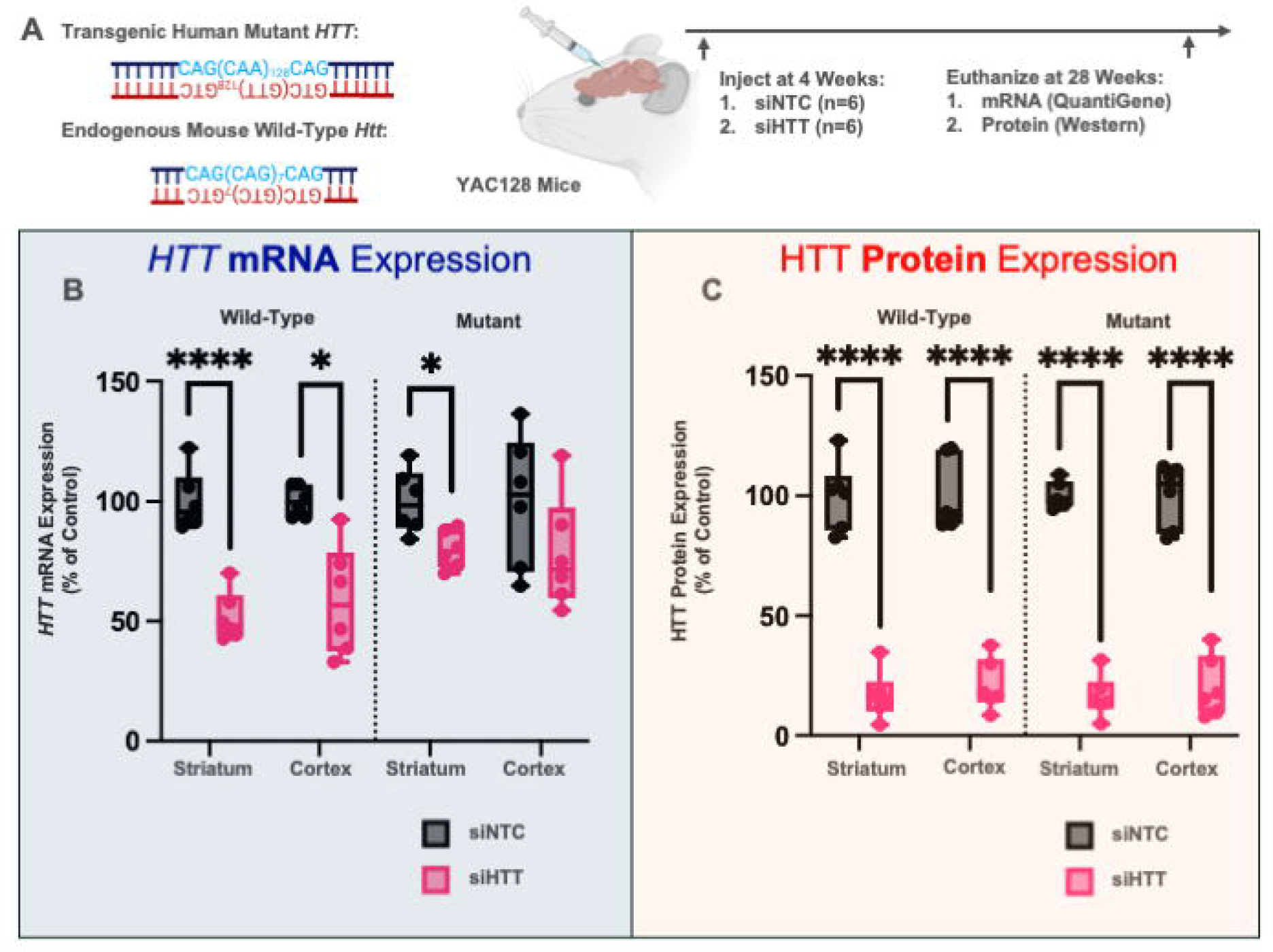
*HTT* mutant and wild-type mRNA but not protein is differentially affected by di-siRNA treatment in YAC128 at 6 months post injection. (A) One month old YAC128 mice were injected ICV with 10 μL of 20 nmol di-siRNA targeting either *HTT* or a non-targeting control (n=6). YAC128 mice express both transgenic human mutant *HTT* with 128 mixed CAA/CAG repeats and endogenous mouse wild-type *Htt* with 7 CAG repeats. Mice were sacrificed six months later at 32 weeks old. After sacrifice striatum and cortical tissue were tested for changes *HTT* mRNA (Quantigene) and protein (automated simple western) expression. (B) Human mutant *HTT* and wild-type mouse *Htt* mRNA levels were measured by Quantigene and normalized to *Hprt* as a housekeeping gene. mRNA levels are displayed as a percentage of their respective non-targeting control value. For comparisons between groups a one-way anova was used with multiple comparisons. For comparisons between siNTC and siHTT treated tissue p = <0.0001 and p= 0.0172 for wild type *Htt* in the striatum and cortex respectively. For mutant *HTT* p = 0.0253 and p = 0.3255 in the striatum and cortex respectively. (C) Human mutant HTT and wild-type mouse Htt protein levels were measured by automated simple western and normalized to vinculin as a loading control. Protein levels are displayed as a percentage of their respective non-targeting control value. For comparisons between groups a one-way anova was used with multiple comparisons. For comparisons between siNTC and siHTT treated tissue, p = <0.0001 and p= <0.0001 for wild type Htt in the striatum and cortex respectively. For mutant HTT p = <0.0001 and p = <0.0001 in the striatum and cortex respectively.

Taken together, our results in BAC-CAG and YAC128 mice suggest that partial mRNA silencing can result in nearly complete silencing at the protein level.

### FISH analysis of di-valent siRNA treated BAC-CAG striatum shows differential nuclear localization of mutant *HTT* and wild-type *Htt* mRNA

We sought to determine whether the discrepancies between wild-type *Htt* and mutant *HTT* mRNA silencing were due to differential localization of mutant mRNA transcripts. We performed FISH on BAC-CAG mouse brain tissue treated with divalent siRNA against *HTT* or with NTC control (samples from study in Figure 1). Samples were cryosectioned, stained, and regionally imaged in the striatum (Figure 3A, 240 μg (10 nmol), one month post injection).

**Figure 3:**
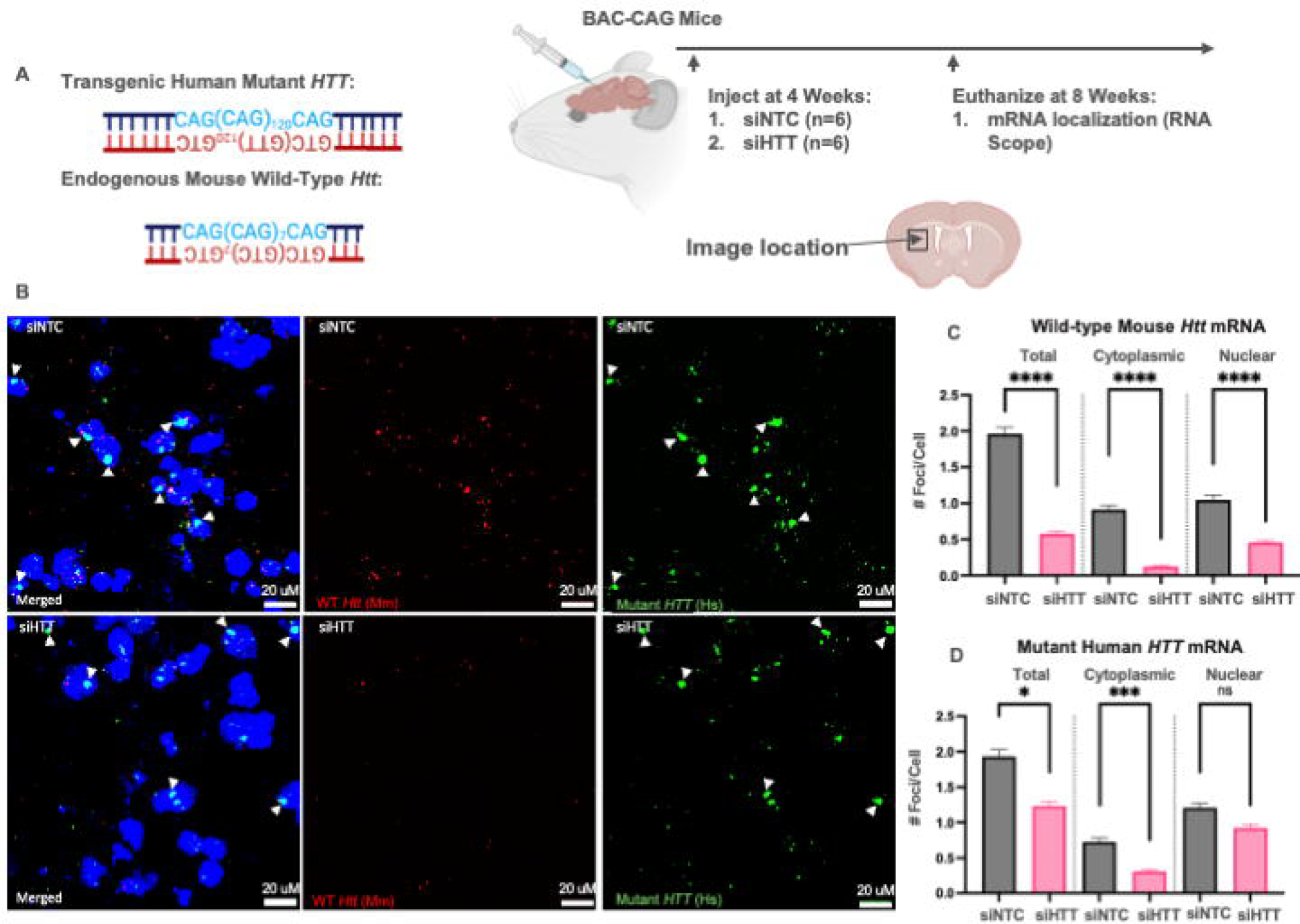
Nuclear mutant but not wild-type huntingtin mRNA fraction is resistant to siRNA-based silencing in striatum of BAC-CAG mice. (A) Three-month-old BAC-CAG mice were injected ICV with 10 μL of 20 nmol di-siRNA targeting either *HTT* or a non-targeting control (n=6). BAC-CAG mice express both transgenic human mutant *HTT* with 120 CAG repeats and endogenous mouse wild-type *Htt* with 7 CAG repeats. Mice were sacrificed one month later at 20 weeks old. After sacrifice striatum and cortical tissue were tested for changes HTT mRNA (Quantigene), protein (automated simple western) expression (figure 1), and mRNA localization (RNA FISH). (B) Representative fluorescent in situ hybridization (FISH) images showing NTC treated (upper) and siHTT (lower) treated BAC-CAG striatum tissue. Probes measure mutant *HTT* (right, green) and wild-type *Htt* (middle, red). Left images show a merge of both probes plus dapi staining cell nuclei (blue). White arrows denote intranuclear clustering. (C-D) Quantification of FISH images (n=3 animals) for mutant *HTT* and wild-type *Htt* in different compartments. All statistical tests were performed using a mixed effects model and analyzed using pairwise comparisons contrasting the impact of siRNA treatment. (B) Counts of *Htt* foci in cellular compartments with siNTC or siHTT treatment (****p<0.0001). (C) Counts of *HTT* foci in cellular compartments with NTC or siHTT treatment (*p=0.04, ***p=0.0001).

Consistent with published observations (7, 8) mutant *HTT* mRNA, but not wild-type *Htt* mRNA, forms large intranuclear clusters (Figure 3B, white arrows). The clusters are defined as foci greater than 1.5 μm^3^ while mRNA foci (volume <1.5 μm^3^) are indicative of single mRNA transcripts. The rest of the distribution was similar except for a slightly higher fraction of individual mutant mRNA foci localized in the nucleus (62% vs 50%) mainly driven by higher number of cytoplasmic foci observed in wild-type *Htt* mRNA.

Using Image J, stereotactically collected striatum images of at least 100 cells per condition were quantified to determine the number of total, nuclear, and cytoplasmic wild-type *Htt* and mutant *HTT* mRNA foci per cell (see methods). For NTC treated brain tissue, there were an average of 1.95 mutant *HTT* vs 2.45 wildtype *Htt* individual foci per cell. Consistent with previously reported data (7), there were high levels of heterogeneity in *HTT* expression, ranging from 0-38 foci for mutant *HTT* and 0-19 foci for wildtype *Htt*. An average of 36% and 26% of cells have no detectable mutant and wild-type huntingtin foci, respectively.

There were 0.72 vs 1.22 cytoplasmic foci/cell for mutant vs wildtype *HTT*. This is consistent with observed lower level of mutant HTT protein, as solely the cytoplasmic mRNA fraction is responsible for HTT protein expression [(6), Supplementary Figure 2].

Di-siRNA treatment significantly reduced wild-type mouse *Htt* mRNA foci in all cellular compartments (p<0.0001, 57% silencing in nucleus; p<0.0001, 87% silencing in cytoplasm; and p<0.0001, 71% silencing in total), with the level of silencing being greater in the cytoplasm than the nucleus (Figure 3B-C). Di-siRNA treatment also significantly reduced the total number of mutant *HTT* mRNA foci (36% silencing, p=0.04) and cytoplasmic foci (57% silencing, p=0.0001), but not nuclear foci (24% silencing, p= 0.21, Figure 3B,D). The observed silencing of cytoplasmic *Htt* and *HTT* mRNA may explain why near-complete silencing of Htt and HTT protein is observed following di-valent siRNA treatment, as only cytoplasmic mRNA is translated into protein. Upon further evaluation of the nuclear fraction, di-siRNA treatment reduced individual mutant *HTT* mRNA foci but had minimal effect on mutant *HTT* mRNA clusters (volume>1.5 μm^3^), consistent with previously reported data for HTT targeting ASO GapmeRs (7).

## Discussion

Here we show that there is difference in the observed level of mutant and wild-type huntingtin mRNA silencing by di-valent siRNA *in vivo* in the brains of two different mouse models of HD. The effect is observed at different ages and various treatment windows. Incomplete mRNA silencing results in near-complete elimination of protein expression. Using advanced FISH, we demonstrate that the discrepancy in mRNA and protein silencing efficiency leads to changes in intracellular mRNA localization due to repeat expansion. Mutant mRNA forms nuclear clusters that are resistant to RNAi, while both nuclear and cytoplasmic wild-type *Htt* mRNA can be silenced. This study is a first to report on the impact of structure of nuclear RNA impacting efficiency of RNAi-based silencing.

It is proven that AGO2, the enzymatic factor driving RNAi activity, can be localized both to the nucleus and cytoplasm, albeit in lesser amounts in the nucleus. Cellular state (ie internal or environmental stressors) have an impact on the relative distribution of AGO2 between two compartments (22). While siRNA can silence RNAs in the nucleus (Figure 3C), the extent to which intracellular localization and clustering impacts siRNA efficacy *in vivo* is not well understood. The relative potency of nuclear RNAi is lower than cytoplasmic RNAi (23). In cells, it is believed that siRNAs are more effective towards cytoplasmic targets, while ASOs are better in targeting nuclear localized transcripts (24). While true in general, we have previously shown the mutant *HTT* nuclear mRNA clusters are highly resistant to GapmeR ASO based silencing (7). Thus nuclear clustering and supramolecular assemblies of mRNA are making them a challenging target for oligonucleotide-based modulation (10, 11).

We observe that *in vivo*, cytoplasmic non-clustered mRNA foci are silenced nearly 90% while silencing of nuclear transcript does not exceed 65%. While it is likely that the data presented supports AGO2-based nuclear silencing, the potential of a redistributive shift in the ratio of nuclear/cytoplasmic mRNA upon siRNA treatment cannot be fully excluded. Recent data from Avidity shows robust siRNA-based silencing of another preferentially nuclear mRNA, DMPK *in vivo*, providing additional indirect evidence for accessibility of the nuclear mRNA to the RNAi modulation (25).

We show that mutant mRNA clusters are resistant toward RNAi based silencing, indicating that the structure and clustering may be a primary factor limiting nuclear RNAi for some targets. Target mRNA secondary structure has been demonstrated to have the potential to impact siRNA efficacy (26). mRNA aggregation is now reported in many cases of repeat expansion disorders (27–29) and is believed to be contributing to the clinical manifestation of disease. In addition, many non-coding RNA targets of interest are nuclear localized and are known to form complex tertiary structures (30). The impact of supramolecular and intermolecular structure on RNAi potency is an important consideration for selection of siRNA targets, especially for non-traditional targets like viral (31) or nuclear clustered repeat-containing transcripts, including *HTT* in HD, and other neurological disorders with nuclear RNA clusters such as C9orf72 in ALS (32).

While the clinical importance of clusters in HTT is not established, it is likely that they may not be readily accessible to oligonucleotides. In this work we report the resistance of these clusters to siRNA-based modulation. Incomplete mRNA silencing translating into potent protein silencing has been reported for other targets and is specifically common in a context of inflammatory involved pathways (33). For silencing by siRNA, nuclear-cytoplasmic distribution of target mRNA may affect efficiency of RNAi based silencing and should be considered when interpreting mRNA silencing data.

## Conclusions

We demonstrate that in multiple mouse models of HD, at two different time points, there is a difference in the observed level of silencing of mutant and wild-type huntingtin mRNA by di-valent siRNA in vivo. The expanded repeats present in the mutant mRNA drive nuclear mRNA clustering, which is resistant to RNAi-based modulation. Despite incomplete silencing at the mRNA level, there is nearly complete silencing of protein expression. Thus, evaluating oligonucleotide efficacy based solely on mRNA level may yield false negative results. Additionally, when higher levels of protein silencing compared to mRNA silencing is observed, the impact of mRNA intracellular localization should be considered.

## Acknowledgements

The authors acknowledge all lab members who kindly reviewed the manuscript and provided editorial suggestions. The authors thank Samuel Hildebrand for his assistance with statistical analysis and Lori Kennington, Vicky Benoit, and Rachael Miller for their assistance in breeding and genotyping mice.

## Authorship Confirmation/Contribution Statement

Conceptualization was performed by S.A., D.O., R.M., and A.K. Visualization was performed by S.A., D.O., A.K., N.A., and M.D. Methodology was designed by S.A., D.O., R.M., and A.K. Investigation was performed by S.A., D.O., R.M, E.S., A.S., J.P., D.E.M., B.B., N.M., and K.Y. Funding acquisition was done by A.K., N.A., M.D., and K.Y. Project administration and supervision were performed by A.K., N.A., and M.D. Writing the original draft was completed by S.A and A.K. Reviewing and editing the original draft was completed by S.A., D.O., A.K., N.A., and M.D.

