## Supplementary figures and images for "mRNA nuclear clustering leads to a difference in mutant huntingtin mRNA and protein silencing by siRNAs *in vivo*"

### Supplementary Figures

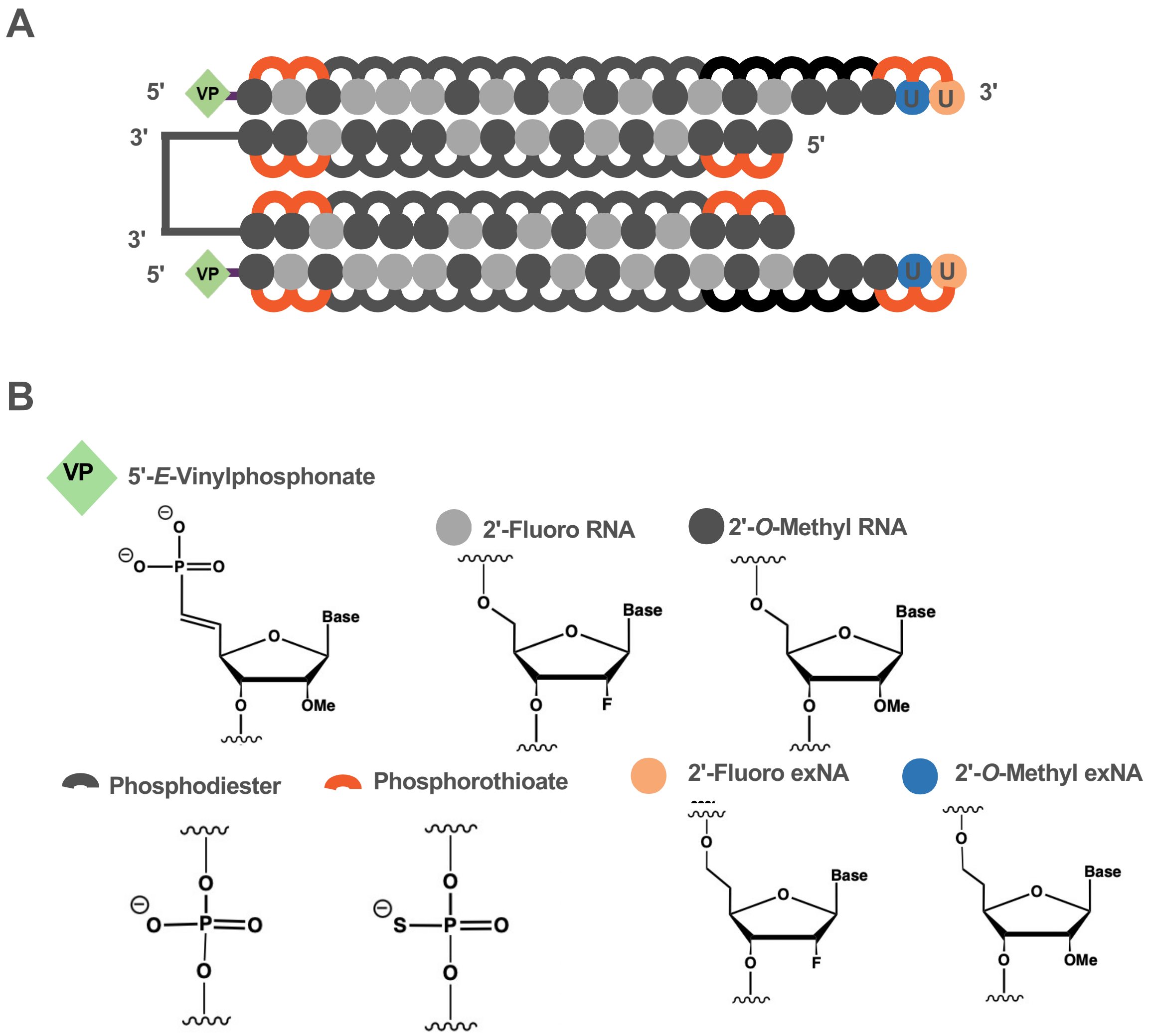

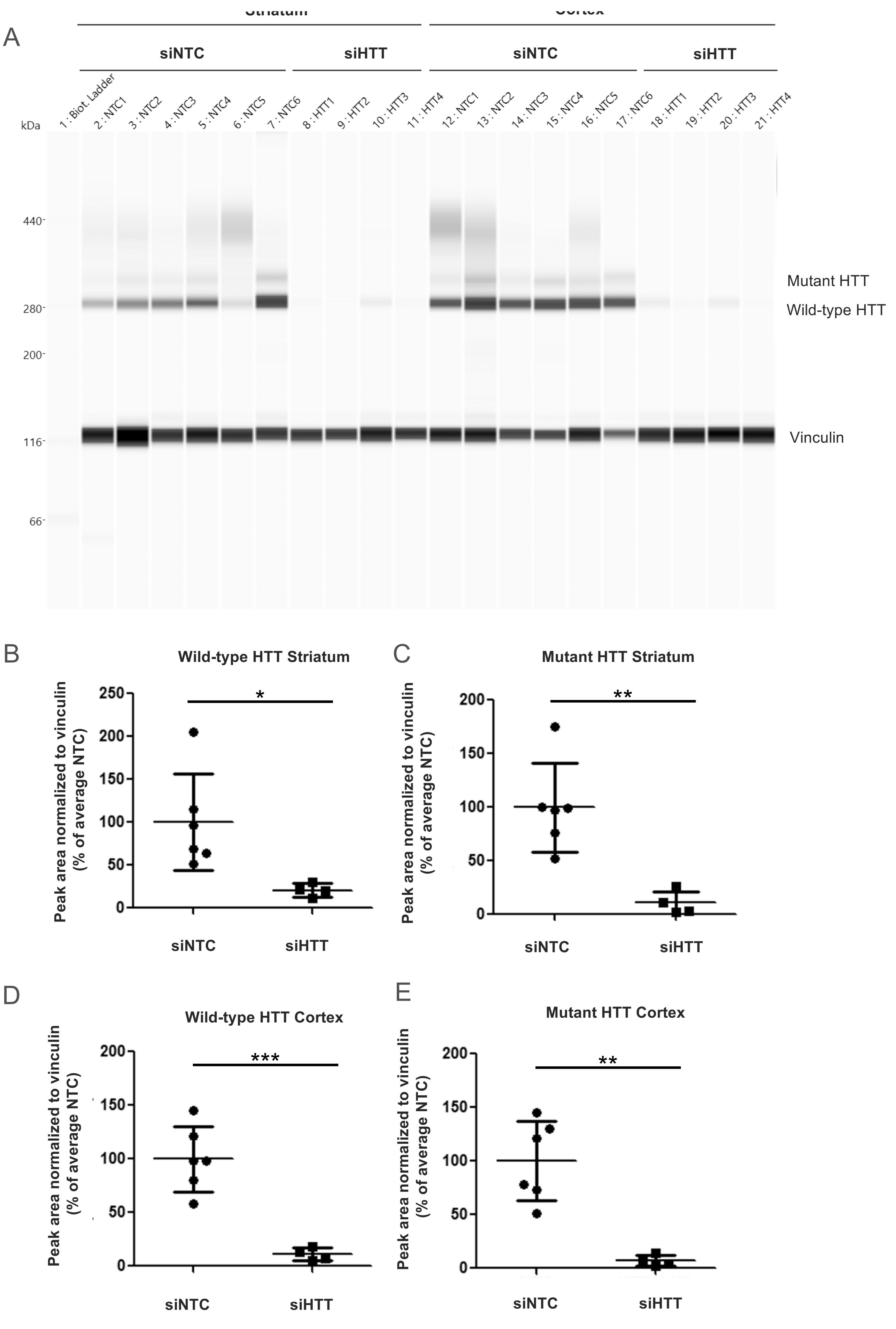

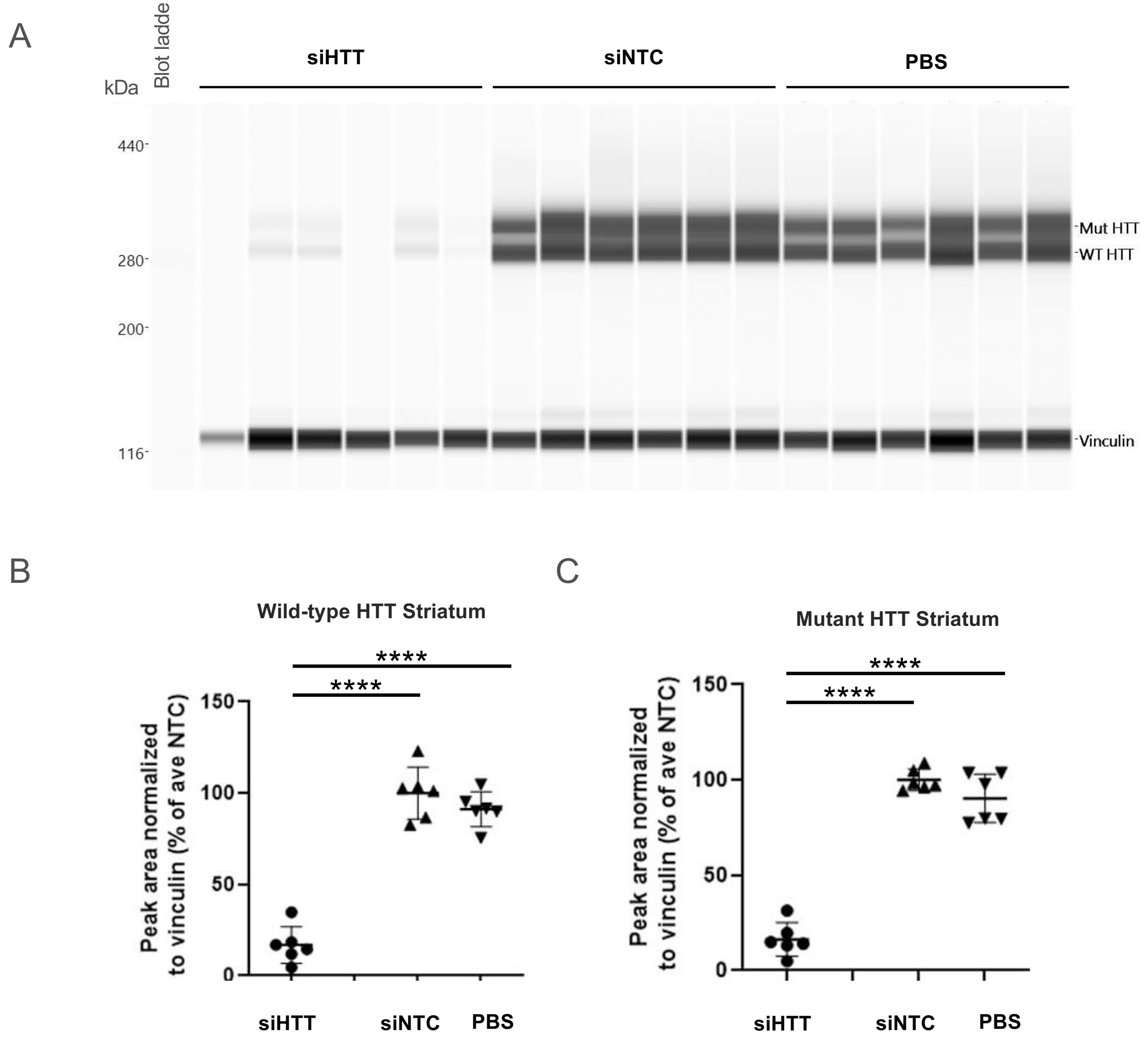

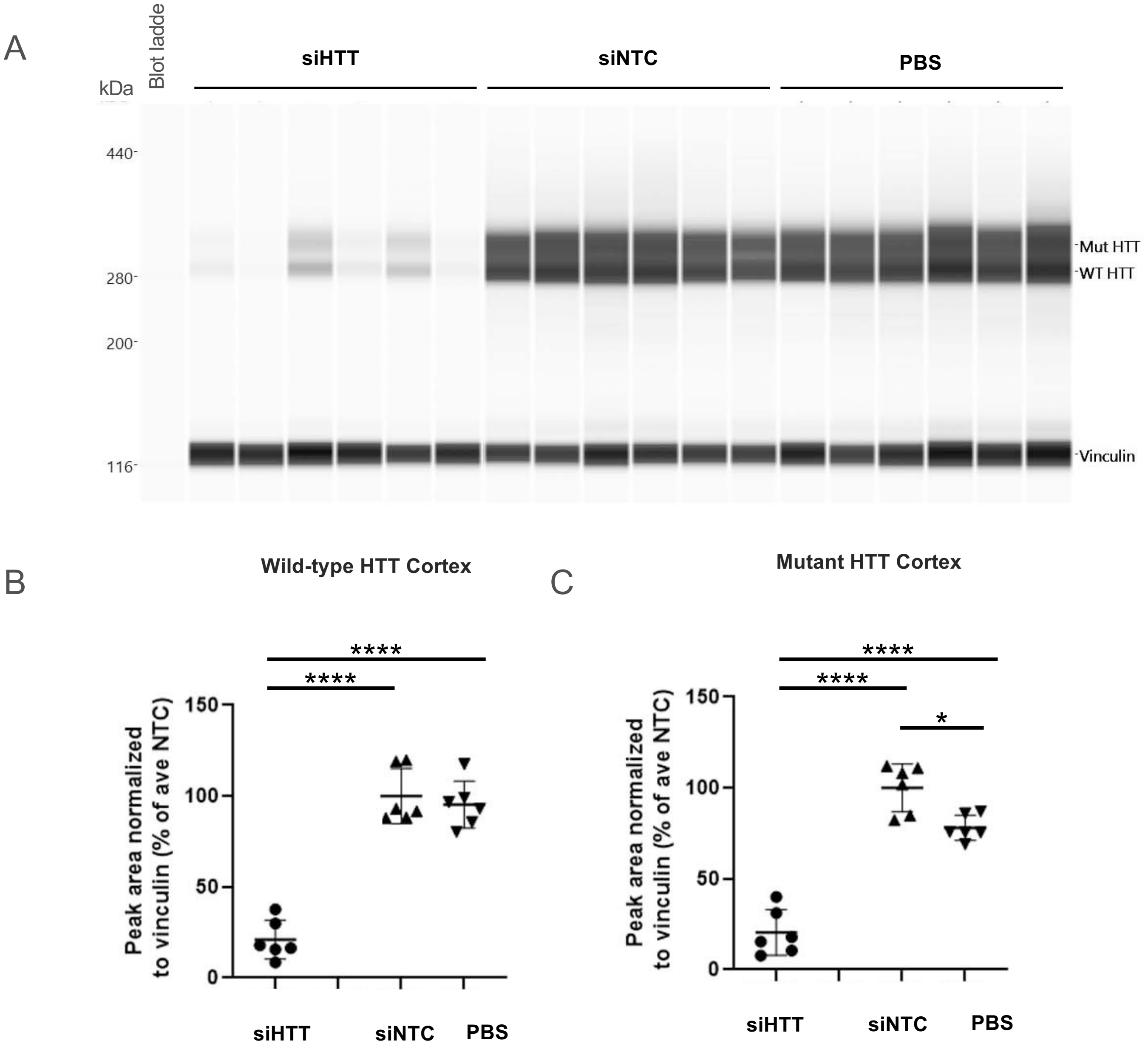
